# New minimally invasive techniques for sample collection in Antarctic seals

**DOI:** 10.1101/2025.07.15.664748

**Authors:** Florencia A. Soto, Habib D. Ahumada, Ángela Vázquez-Calvo, Antonio Alcamí, Javier Negrete

## Abstract

Remote or minimally invasive sampling techniques are essential in wildlife research, particularly for Antarctic seals. These methods reduce handling time, minimize animal stress, and improve biosecurity by preventing pathogen transmission, especially relevant during the current outbreak of highly pathogenic avian influenza affecting Antarctic megafauna. Research on ice-dependent seals is logistically challenging and often requires chemical immobilization, which is costly, time-consuming, and poses risks to both animals and researchers. To overcome these limitations, we developed a simple, low-cost remote sampling device to collect nasal swabs from seals without physical contact. The device consists of a telescopic pole with a swab attached to its end. Researchers approached seals resting on ice floes or land using inflatable boats, without disembarking, thus minimizing field exposure. Over ten days, we collected ten nasal swabs from leopard seals (*Hydrurga leptonyx*, n=4), Weddell seals (*Leptonychotes weddellii*, n=4), and southern elephant seals (*Mirounga leonina*, n=2). All samples tested negative for avian influenza virus (INF-A) using RT-qPCR. This methodology is adaptable to various species and conditions, requires minimal equipment, and can be implemented by small teams. It provides a safe, efficient tool for disease surveillance in remote polar environments, advancing marine mammal research while minimizing disturbance to wildlife.

## 1 INTRODUCTION

The primary advantage of non-invasive or minimally invasive techniques (MIT) lies in its ability to obtain biological material without causing harm to the target organism, thereby improving animal welfare during fieldwork (Fietz et al., 2016; Taberlet et al., 1999; Van Cise et al., 2024). Those methods include the use of long-handled tools to collect samples such as nasal mucus, feces, hair, or sloughed skin, without the need for direct handling or sedation. These approaches are particularly valuable in light of the increasing risk of zoonotic disease transmission and the emergence of novel pathogens. In addition, non-invasive sampling techniques minimize the risk of mortality associated with sedation, as documented in leopard seals (*Hydrurga leptonyx*) and Weddell seals (*Leptonychotes weddellii*), where anesthesia has resulted in fatal overdoses or complications from underlying health conditions (Higgins et al., 2002; Shero et al., 2025).

Research on Antarctic pinnipeds is crucial for understanding the health of ecosystems and the emergence of wildlife diseases. As apex and mesopredator species, they play a key role in the Antarctic food web. Their health reflects environmental changes, making them effective sentinels for monitoring the status of the ecosystem (Cappozzo et al., 2013; Reddy, 2001). Due to the logistical challenges and ethical considerations involved in handling these animals, we are emphasizing the importance of minimally invasive research techniques. These techniques allow health indicators to be monitored without causing significant stress to the animals, thereby providing valuable data for conservation and management efforts.

A variety of non-invasive sampling tools have been developed and successfully applied in pinniped research over the past decades. For instance, Gemmell & Majluf (1997), introduced a projectile biopsy technique to collect tissue samples from South American fur seals (*Arctocephalus australis*) with minimal disturbance. Swanson et al. (2006), proposed collecting naturally sloughed skin cells from ringed seals (*Pusa hispida*) for DNA analysis. More recently, Peralta et al. (2020) designed a tool using a 3.5-m aluminium pole and adhesive-coated tubes to noninvasively collect hair samples on South American sea lion (*Otaria byronia*). Rivera-Luna et al. (2023) applied a telescopic device fitted with Petri dishes to remotely collect nasal mucus expectorations from free-ranging sea lions (*O. byronia*), enabling detection of halarachnid mite infestations without the need for animal handling.

These methodological advances are especially relevant in the current context of the highly pathogenic avian influenza (HPAI) panzootic, which has caused the death of over 50,000 pinnipeds in South America (Kuiken et al., 2025; Pardo-Roa et al., 2025; Uhart et al., 2024; Ulloa et al., 2023). The recent spread of HPAI H5 to the Antarctic region has resulted in multiple mortality events among seabirds and marine mammals (Banyard et al., 2024), with confirmed cases being reported on the Antarctic Peninsula by February 2024 (*Scientific Committee on Antarctic Research. 2024*). In this urgent scenario, the application of MIT-based strategies becomes critical for monitoring pinniped populations and detecting emerging pathogens. Therefore, in this study, we present a practical, field-tested technique for remotely collecting biological samples from seals that is both minimally invasive and significantly reduces the need for physical handling and field exposure.

## 2 METHODS

### 2.1 Study Area

This study was conducted over ten consecutive days, from 13th to 22nd January 2024, during the Austral summer, in the Antarctic Specially Protected Area (ASPA) N° 134 (64°09’S, 60°57’W), in the northern sector of the Danco Coast, Antarctic Peninsula. The species sampled were Weddell seals (*Leptonychotes weddelli*), leopard seals (*Hydrurga leptonyx*) and Southern elephant seal (*Mirounga leonina*). The seals were located on small ice floes that could be accessed by inflatable boat, or on land that could be accessed on foot.

### 2.2 Sampling device and team roles

An aluminium telescopic pole (2.5 m), metal clamp and swab were used to collect nasal swabs remotely. A three-person field team was enough (Fig.1. 1):

- Operator (sampler): handles the telescopic pole and performs the swabbing.
- Assistant: monitors the animal’s behavior, times the procedure, and manages the sampling equipment and sample storage.
- Sailor: monitors the movement of surrounding ice and ensures the safety of the team and vessel during the procedure.

**Figure 1.**
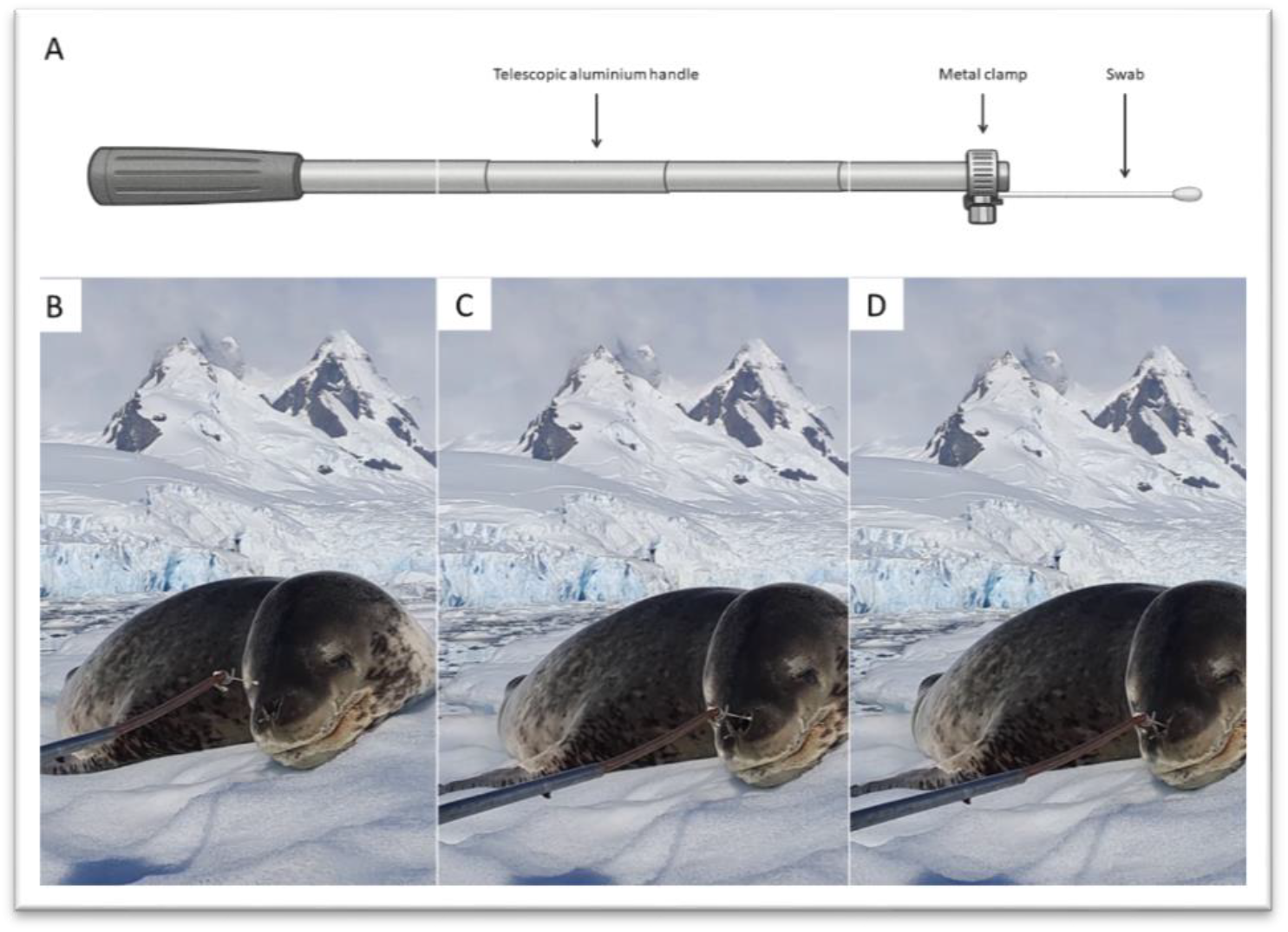
A) Diagram of the sampling device, which consists of a telescopic aluminium handle, a metal clamp and a swab. B) The seal rests on an ice floe while the pole is slowly and carefully approached. C) The swab is held close to the animal’s nostril, waiting for the seal to exhale. As the seal exhales, the nostril opens to allow the swab to enter. D) The swab remains briefly in the nasal cavity during inhalation and is then removed and placed in a vial containing RNA/DNA Shield. Photo: Florencia Soto.

All team members wore biosafety gear, including disposable gloves, safety goggles, and face masks, in accordance with strict biosafety protocols and prior to each sampling event the telescopic pole was disinfected using a quaternary ammonium compound.

### 2.3 Sampling procedure

Each sampling event was carried out in a quiet and controlled environment to reduce stress on the animal, boat engines were turned off and radios silenced before approaching. All procedures were completed within 2–5 minutes per individual.

The procedure consisted of the following steps:

#### 1. Approach (Figure 1.2A)

- When the animal was located on an ice floe, the sailor approached using the boat engine up to approximately 3 meters, then turned off the engine and paddled slowly to bring the vessel within sampling range.
- When the seal was on the beach, the team approached on foot, walking slowly and cautiously to avoid disturbing the animal.

In both cases, the operator extended the telescopic pole toward the seal, carefully aligning the swab tip with one of the animal’s nostrils. Movements were slow and deliberate throughout to minimize the risk of startling the animal.

#### 2. Positioning (Figure 1.2B)

The swab was held just in front of the nostril, waiting for a natural exhalation. This brief exhalation causes the nostril to open, facilitating swab insertion.

#### 3. Sampling (Figure 1.2C)

During the following inhalation, the swab was gently inserted into the nasal cavity and rotated for 5–10 seconds against the mucous membranes to collect the sample. The swab was then carefully withdrawn.

#### 4. Sample Preservation

The swab was immediately placed into a sterile vial containing RNA/DNA Shield to stabilize nucleic acids for subsequent molecular analysis. The vial was securely sealed and labeled.

The telescopic pole was thoroughly disinfected using a quaternary ammonium solution after each individual, ensuring coverage of the swab attachment point and any part that may have come into contact with the animal or environment. Gloves were changed between individuals to prevent cross-contamination.

### 2.3 Viral RNA extraction and influenza A detection by RT-qPCR

The molecular diagnostic of INF-A was carried out in the laboratory of the Spanish Antarctic Base Gabriel de Castilla (Deception Island, South Shetland Islands, Antarctica). Procedures for viral RNA extraction and RT-qPCR were as described previously (18). Brieftly, RNA was extracted using the Maxwell RSC Viral Total Nucleic Acid Purification kit in a Maxwell® RSC 48 Instrument (Promega). A blank sample was included in each extraction batch that was used as a negative control for RT-qPCRs. Before extraction, around 500 ng of mouse RNA from L929 cells was added to each sample as carrier for nucleic acid isolation. This carrier was later used as extraction RT-qPCR control (see more details in (Aguado et al., 2024). The presence of INF-A in the samples was determined by RT-qPCR in a CFX Opus 96 Real-Time PCR System using One-Step Multiplex RT-qPCR Supermix (BIO-RAD), specific primers Inf-A Fw1: CAAGACCAATCYTGTCACCTCTGAC;Inf-A Rv1: GCATTYTGGACAAAVCGTCTACG; Inf-A Fw2: CAAGACCAATYCTGTCACCTYTGAC and Inf-A Rv2: GCATTTTGGATAAAGCGTCTACG) and a fluorescent labelled probed (Inf-A Probe: /56-ROXN/TGCAGTCCTCGCTCACTGGGCACG/3IAbRQSp/) against a partial sequence from segment 7 (M1) of INF-A. A negative extraction control, non-template control and a positive control were included in each plate. As extraction control, partial hypoxanthine guanine phosphoribosyltransferas (HPRT) gene from mouse was targeted and detected in each sample using specific primers (Fw: TTCTTTGCTGACCTGCTGGA; Rv: ACAATCAAGACATTCTTTCCAGTT) and fluorescent-labelled probe (/5HEX/ATTGGTGGA/ZEN/GATGATCTCTCAACT/3IABkFQ/). Samples with a cycle threshold (Ct) value lower than 40 were considered positive by RTqPCR. Results were analysed using CFX Maestro 2.3 software (BIO-RAD).

## RESULTS

We attempted to sample 13 seals, successfully collecting 10 nasal swabs without any direct contact with the animals. Sampling attempts failed on two crabeater seals, which were highly alert to our presence, and one Weddell seal, which woke up suddenly during radio communication, failed. Information on each seal, including the date, location, species, age group, sex, and number of attempts, can be found in Table 1.

**Table 1.** Sampling details of Antarctic seals, including date, location, species, age class, sex and attempts.

| Date | Location | Species | Age Class | Sex | Attempts | RTq PCR results |  |  |
| --- | --- | --- | --- | --- | --- | --- | --- | --- |
|  |  |  |  |  |  | Extraction control | INF-A | Result |

|  |  |  |  |  |  | (CT value) | (CT value) |  |
| --- | --- | --- | --- | --- | --- | --- | --- | --- |
| 13 Jan 2024 | Land | *L. weddellii* | Adult | Male | Success | 29.6 | NA | Negative |
| 13 Jan 2024 | Land | *M. leonina* | Juvenile | Female | Success | 35.6 | NA | Negative |
| 13 Jan 2024 | Ice | *L. carcinophaga* | Juvenile | Undetermined | Failure | - | - | - |
| 16 Jan 2024 | Ice | *L. weddellii* | Adult | Female | Success | 35.9 | NA | Negative |
| 16 Jan 2024 | Ice | *L. weddellii* | Adult | Male | Success | 35.7 | NA | Negative |
| 16 Jan 2024 | Ice | *L. weddellii* | Adult | Male | Success | 35.0 | NA | Negative |
| 16 Jan 2024 | Land | *M. leonina* | Juvenile | Undetermined | Success | 37.9 | NA | Negative |
| 21 Jan 2024 | Ice | *L. weddellii* | Weddell | Male | Failure | - | - | - |
| 21 Jan 2024 | Ice | *H. leptonyx* | Adult | Male | Success | 34.5 | NA | Negative |
| 21 Jan 2024 | Ice | *L. carcinophaga* | Adult | Undetermined | Failure | - | - | - |
| 22 Jan 2024 | Ice | *H. leptonyx* | Adult | Male | Success | 35.8 | NA | Negative |
| 22 Jan 2024 | Ice | *H. leptonyx* | Adult | Male | Success | 34.5 | NA | Negative |
| 22 Jan 2024 | Ice | *H. leptonyx* | Adult | Male | Success | 34.0 | NA | Negative |

INF-A was not detected in the nasal swabs, and consequently, H5 testing was not performed.

## DISCUSSION

In recent decades, various minimally invasive sampling techniques have been developed to minimize the risks and logistical challenges associated with the chemical immobilization of pinnipeds, particularly in remote polar regions. These methods reduce the likelihood of disease transmission between animals by using sterile, single-use materials, while also enhancing researcher safety by minimizing the risk of zoonotic exposure. This approach is especially relevant in the current context of emerging infectious diseases in wildlife and the growing need for stringent biosecurity measures in polar research.

In this framework, and despite the limited sample size in the present study, we aimed to introduce an innovative sampling methodology tailored to hard-to-access species. The procedure can be implemented by small field teams using minimal and readily available equipment, and can be adjusted to suit the behavior of each seal species. A key practical insight was that sampling success improved when external noise was reduced, specifically by switching off engines and silencing radios, to avoid disturbing the animals. Notably, the sampling pole is inexpensive, easy to construct, and simple to operate, requiring no specialized training, in contrast to UAVs or other remote-controlled technologies, which often involve higher costs, regulatory constraints, and the need for trained personnel (Goebel et al., 2015; Hodgson et al., 2013; Koski et al., 2009).

Although the animals often reacted to the initial approach, they generally remained calm and undisturbed throughout the sampling process. However, crabeater seals were an exception, showing heightened sensitivity to human presence and remaining alert even when approached cautiously. Based on these observations, we suggest that future efforts could explore the use of natural ice formations on large icebergs as visual barriers, allowing the operator to extend the pole and collect samples without being fully visible to the animal.

Building on these findings, our method offers a practical alternative to traditional immobilization techniques by significantly reducing animal stress, field exposure, and biosecurity risks, while maintaining sample integrity. In our case, all samples tested negative for INF-A; therefore, no additional testing for H5 was conducted. Performing the RT-qPCR tests in a laboratory set up in the Spanish Antarctic Base Gabriel de Castilla (Deception Island) allowed the rapid diagnostic of the samples in Antarctica, avoiding the need to send the samples back to the home laboratories. Nonetheless, given the confirmed spread of highly pathogenic avian influenza (HPAI) to Antarctica (Banyard et al., 2024; Kuiken et al., 2025), including reported infections in pinnipeds (18, MCIU press release), the implementation of this method offers a valuable tool for disease surveillance. Its ease of use and minimal invasiveness make it suitable not only for Antarctic pinnipeds but also for other species affected by HPAI (Alava et al., 2024; Gadzhiev et al., 2024; Lair et al., 2024; Rimondi et al., 2024; Szteren & Franco-Trecu, 2024; Uhart et al., 2024) or other viral pathogens.

Ultimately, our approach contributes to improved disease monitoring in marine mammals while promoting animal welfare and field safety. Its adaptability, low cost, and minimal invasiveness make it suitable not only for Antarctic pinnipeds but potentially for other wild species affected by viral pathogens in remote regions.

## Acknowledgments

The authors support and advocate for the Argentinian Scientific Program, understanding science as an act of sovereignty. The RT-qPCR analysis was funded by Consejo Superior de Investigaciones Científicas (CSIC) (Grant 202320E224) and supported by the Spanish Polar Program. Fieldwork was conducted under Permit No. [2023-FEAMB-CT-GA-49], issued by the Dirección Nacional del Antártico-Instituto Antártico Argentino (DNA-IAA). We want to thank those involved for fieldwork assistance in Antarctica; especially we thank Mauro Rozas Sia, and Rosana Sandler, Marianela Beltán and Emiliano Martinez for their invaluable assistance during fieldwork. Finally, we deeply acknowledge the constructive comments and valuable suggestions made by Ralph E.T. Vanstreels.

## Author Contributions

Conceptualization: F.A.S., H.D.A., J.N. Data curation: F.A.S., H.D.A. Formal analysis: F.A.S., H.D.A., J.N., A.V.C; Funding acquisition: J.N., A.A.; Investigation: F.A.S., H.D.A., A.V.C., A.A., J.N. Methodology: F.A.S., H.D.A., J.N. Validation: F.A.S., H.D.A., J.N., A.V.C., A.A. Writing-original draft: F.A.S., H.D.A., A.V.C., A.A., J.N. Writing review and editing: F.A.S., H.D.A., J.N. All authors have read and agreed to the published version of the manuscript.

## Notes

### Competing Interest Statement

The authors have declared no competing interest.

## REFERENCES

Aguado, B., Begeman, L., Günther, A., Iervolino, M., Soto, F., Vanstreels, R. E., Reade, A., Coerper, A., Wallis, B., & Alcamí, A. (2024). Searching for high pathogenicity avian influenza virus in Antarctica. Nature Microbiology, 1–3. 10.1038/s41564-024-01868-7.

Alava, J. J., Tirapé, A., Denkinger, J., Calle, P., Rosero R. P., Salazar, S., Fair, P. A., & Raverty, S. (2024). Endangered Galápagos sea lions and fur seals under the siege of lethal avian flu: A cautionary note on emerging infectious viruses in endemic pinnipeds of the Galápagos Islands. Frontiers in Veterinary Science, 11, 1457035. 10.3389/fvets.2024.1457035

Banyard, A. C., Bennison, A., Byrne, A. M., Reid, S. M., Lynton-Jenkins, J. G., Mollett, B., De Silva, D., Peers-Dent, J., Finlayson, K., & Hall, R. (2024). Detection and spread of high pathogenicity avian influenza virus H5N1 in the Antarctic Region. Nature Communications, 15(1), 7433. 10.1038/s41467-024-51490-8

Cappozzo, H. L., Panebianco, M., & Túnez, J. I. (2013). Marine mammals in a changing world. Marine Ecology in a Changing World. CRC Press, Boca Ratón, Florida, Estados Unidos, 219–259.

Fietz, K., Galatius, A., Teilmann, J., Dietz, R., Frie, A. K., Klimova, A., Palsbøll, P. J., Jensen, L. F., Graves, J. A., & Hoffman, J. I. (2016). Shift of grey seal subspecies boundaries in response to climate, culling and conservation. Molecular Ecology, 25(17), 4097–4112. 10.1111/mec.13748

Gadzhiev, A., Petherbridge, G., Sharshov, K., Sobolev, I., Alekseev, A., Gulyaeva, M., Litvinov, K., Boltunov, I., Teymurov, A., Zhigalin, A., Daudova, M., & Shestopalov, A. (2024). Pinnipeds and avian influenza: A global timeline and review of research on the impact of highly pathogenic avian influenza on pinniped populations with particular reference to the endangered Caspian seal (Pusa caspica). Frontiers in Cellular and Infection Microbiology, 14, 1325977. 10.3389/fcimb.2024.1325977

Gemmell, N. J., & Majluf, P. (1997). Projectile biopsy sampling of fur seals. Marine Mammal Science, 13(3), 512–516.

Goebel, M. E., Perryman, W. L., Hinke, J. T., Krause, D. J., Hann, N. A., Gardner, S., & LeRoi, D. J. (2015). A small unmanned aerial system for estimating abundance and size of Antarctic predators. Polar Biology, 38(5), 619–630. 10.1007/s00300-014-1625-4

Higgins, D. P., Rogers, T. L., Irvine, A. D., & Hall-Aspland, S. A. (2002). Use of midazolam/pethidine and tiletamine/zolazepam combinations for the chemical restraint of leopard seals (Hydrurga leptonyx). Marine Mammal Science, 18(2), 483–499. 10.1111/j.1748-7692.2002.tb01050.x

Hodgson, A., Kelly, N., & Peel, D. (2013). Unmanned Aerial Vehicles (UAVs) for Surveying Marine Fauna: A Dugong Case Study. PLoS ONE, 8(11), e79556. 10.1371/journal.pone.0079556

Koski, W. R., Allen, T., Ireland, D., Buck, G., Smith, P. R., Macrander, A. M., Halick, M. A., Rushing, C., Sliwa, D. J., & McDonald, T. L. (2009). Evaluation of an Unmanned Airborne System for Monitoring Marine Mammals. Aquatic Mammals, 35(3), 347–357. 10.1578/AM.35.3.2009.347

Kuiken, T., Vanstreels, R. E. T., Banyard, A., Begeman, L., Breed, A., Dewar, M., Fijn, R., Serafini, P. P., Uhart, M., & Wille, M. (2025). tEmergence, spread, and impact of high-pathogenicity avian influenza H5 in wild birds and mammals of South America and Antarctica. Conservation Biology, e70052. 10.1111/cobi.70052

Lair, S., Quesnel, L., Signore, A. V., Delnatte, P., Embury-Hyatt, C., Nadeau, M.-S., Lung, O., Ferrell, S. T., Michaud, R., & Berhane, Y. (2024). Outbreak of Highly Pathogenic Avian Influenza A(H5N1) Virus in Seals, St. Lawrence Estuary, Quebec, Canada1. Emerging Infectious Diseases, 30(6). 10.3201/eid3006.231033

MCIU (Ministerio de Ciencia, Innovación y Universidades). (2024, julio 14). https://www.ciencia.gob.es/en/Noticias/2024/Julio/gripe-aviar-antartida.html

Pardo-Roa, C., Nelson, M. I., Ariyama, N., Aguayo, C., Almonacid, L. I., Gonzalez-Reiche, A. S., Muñoz, G., Ulloa, M., Ávila, C., & Navarro, C. (2025). Cross-species and mammal-to-mammal transmission of clade 2.3. 4.4 b highly pathogenic avian influenza A/H5N1 with PB2 adaptations. Nature Communications, 16(1), 2232.

Peralta, D. M., Ibañez, E. A., Lucero, S., Cappozzo, H. L., & Túnez, J. I. (2020). A new minimally invasive and inexpensive sampling method for genetic studies in pinnipeds. Mammal Research, 65(1), 11–18. 10.1007/s13364-019-00453-2

Reddy, M. (2001). Marine Mammals as Sentinels of Ocean Health in CRC Handbook of Marine Mammal Medicine, Dierauf, LA, Gulland, FMD, Eds.

Rimondi, A., Vanstreels, R. E. T., Olivera, V., Donini, A., Lauriente, M. M., & Uhart, M. M. (2024). Highly Pathogenic Avian Influenza A(H5N1) Viruses from Multispecies Outbreak, Argentina, August 2023. Emerging Infectious Diseases, 30(4). 10.3201/eid3004.231725

Rivera-Luna, H., Kniha, E., Muñoz, P., Painean, J., Balfanz, F., Hering-Hagenbeck, S., Prosl, H., Walochnik, J., Taubert, A., & Hermosilla, C. (2023). Non-invasive detection of Orthohalarachne attenuata (Banks, 1910) and Orthohalarachne diminuata (Doetschman, 1944) (Acari: Halarachnidae) in free-ranging synanthropic South American sea lions Otaria flavescens (Shaw, 1800). International Journal for Parasitology: Parasites and Wildlife, 21, 192–200. 10.1016/j.ijppaw.2023.06.001

Scientific Committee on Antarctic Research. (2025). Sub-Antarctic and Antarctic highly pathogenic avian influenza H5N1 monitoring project. Retrieved from https://scar.org/library-data/avian-flu#cases. (s. f.).

Shero, M. R., Burek-Huntington, K., McCorkell, R., Nadler, S. A., Rzucidlo, C. L., Klink, A. C., Hindle, A. G., Burns, J. M., & Johnson, S. (2025). Novel presentation and pathophysiology of heavy parasitic burdens in Weddell seals (Leptonychotes weddellii) during sedation. BMC Veterinary Research, 21(1), 300. 10.1186/s12917-025-04740-w

Swanson, B., Kelly, B., Maddox, C., & Moran, J. (2006). Shed skin as a source of DNA for genotyping seals. Molecular Ecology Notes, 6(4), 1006–1009. 10.1111/j.1471-8286.2006.01473.x

Szteren, D., & Franco-Trecu, V. (2024). Incidence of highly pathogenic avian influenza H5N1 in pinnipeds in Uruguay. Diseases of Aquatic Organisms, 160, 65–74. 10.3354/dao03827

Taberlet, P., Waits, L. P., & Luikart, G. (1999). Noninvasive genetic sampling: Look before you leap. Trends in Ecology & Evolution, 14(8), 323–327. 10.1016/s0169-5347(99)01637-7

Uhart, M. M., Vanstreels, R. E., Nelson, M. I., Olivera, V., Campagna, J., Zavattieri, V., Lemey, P., Campagna, C., Falabella, V., & Rimondi, A. (2024). Epidemiological data of an influenza A/H5N1 outbreak in elephant seals in Argentina indicates mammal-to-mammal transmission. Nature Communications, 15(1), 9516. 10.1038/s41467-024-53766-5

Ulloa, M., Fernández, A., Ariyama, N., Colom-Rivero, A., Rivera, C., Nuñez, P., Sanhueza, P., Johow, M., Araya, H., & Torres, J. C. (2023). Mass mortality event in South American sea lions (Otaria flavescens) correlated to highly pathogenic avian influenza (HPAI) H5N1 outbreak in Chile. Veterinary Quarterly, 43(1), 1–10. 10.1080/01652176.2023.2265173

Van Cise, A. M., Switzer, A. D., Apprill, A., Champagne, C. D., Chittaro, P. M., Dudek, N. K., Gavery, M. R., Hancock-Hanser, B. L., Harmon, A. C., & Jaffe, A. L. (2024). Best practices for collecting and preserving marine mammal biological samples in the ‘omics era. Marine Mammal Science, 40(4), e13148. 10.1111/mms.13148

